# Acetobase: database update and reanalysis of formyltetrahydrofolate synthetase amplicon sequencing data from the anaerobic digesters

**DOI:** 10.1101/2021.09.19.460943

**Authors:** Abhijeet Singh, Anna Schnürer

**Author notes:** For correspondence.; Address: Department of Molecular Sciences, Box 7025, 75007 Uppsala, Sweden.

## Abstract

AcetoBase is a public repository and database published in 2019, for the formyltetrahydrofolate synthetase (FTHFS) sequences. It is the first systematic collection of bacterial formyltetrahydrofolate nucleotide and protein sequences from the genomes and metagenome assembled genomes (MAGs), as well as sequences generated by clone library sequencing. In addition, AcetoBase was first to establish connection between FTHFS gene with the Wood-Ljungdahl pathway and 16S rRNA genes. Since the publication of AcetoBase, significant improvements were seen in the taxonomy of many bacterial lineages and accessibility/availability of public genomics and metagenomics data. Thus, an update to the AcetoBase database with new sequence data and taxonomy has been made along with improvements in web-functionality and user interface. The update in AcetoBase reference database version 2 was furthermore evaluated by reanalysis of publicly accessible FTHFS amplicon sequencing data previously analysed with AcetoBase version 1. The latest database update showed significant improvements in the taxonomic assignments of FTHFS sequences. AcetoBase with its enhancements in functionality and content is publicly accessible at https://acetobase.molbio.slu.se.

## INTRODUCTION

Formyltetrahydrofolate synthetase (FTHFS) is a key marker gene of the Wood-Ljungdahl pathway (WLP) of acetogenesis (Lovell, 1994; Lovell and Leaphart, 2005). In the enzymatic process of acetogenesis, FTHFS facilitates the ATP dependent conversion of formate to formyl-tetrahydrofolate in the methyl branch of WLP. Although FTHFS gene is also present in other bacteria *viz*. syntrophic acid oxidising bacteria, sulphate reducing bacteria, methanogens *etc*., only acetogens use it in the true sense of acetogenesis (Drake, 1994; Lovell, 1994; Singh et al., 2019, 2020). For the extensive culture independent investigations into the ecology of acetogenic communities in different environments, modern molecular methods are of immense importance (Lovell, 1994). In this context, FTHFS has been used for decades as a molecular marker for the identification of potential acetogenic bacterial communities (*e*.*g*. Lovell and Hui, 1991; Lovell, 1994; Leaphart and Lovell, 2001; Leaphart et al., 2003). As discussed earlier in several studies, acetogens are phylogenetically diverse, metabolically very agile and are important in many environments (*e*.*g*. Drake, 1994; Drake et al., 2008, 2013; Singh et al., 2019). Recently, a high-throughput sequencing method was set-up targeting the FTHFS gene and analysis of different anaerobic digesters revealed the structure and temporal dynamics of potential acetogenic bacterial communities using a metagenomics approach (Singh et al., 2020, 2021b; Singh, 2021).

The analysis of any high-throughput amplicon sequencing data highly relies on the quality and taxonomic accuracy of reference database. For the reliable identification and annotation of high-throughput amplicon sequencing data, we have created and published AcetoBase as a first curated database and repository of FTHFS sequences (Singh et al., 2019). Since its initial publication, there have been a significant increase in the number of genomic and metagenome assembled genomes (MAGs) datasets, deposited in public databases, which contain FTHFS sequences. Additionally, under the influence of recent technological improvements in genomics and bioinformatics, the taxonomy of many bacterial lineages has been redefined. Thus, it is relevant and necessary to update AcetoBase with recent and updated information for the correct identification and annotation of FTHFS sequences. In this paper, we presented the recent changes and updates made in AcetoBase and results from a reanalysis of the FTHFS sequence data generated in our previous studies.

## NEW DEVELOPMENTS

### Database content

AcetoBase version 2 contains around 22,300 protein and 17,000 nucleotide sequences retrieved from public repositories. These sequences belong to 8,439 distinct taxonomic identifiers. The version 1 of AcetoBase contained approximately 18,000, 13,000 and 7,928 protein sequences, nucleotide sequences and taxonomic identifiers. Thus, the new update increased the AcetoBase sequence (protein and nucleotide) and taxonomic identifier diversity by about 4,300, 4,000 and 511, respectively. The major phyla associated with AcetoBase version 2 sequences are Firmicutes, Actinobacteria, Proteobacteria and Bacteroidetes (**Fig. 1**). An interactive version of **Fig. 1** is available at https://acetobase.molbio.slu.se/home. Most sequences (nucleotide and protein) in AcetoBase version 1 were from isolated or characterised bacteria, however, in version two, in addition to sequences from bacterial isolates, FTHFS sequences from metagenome assembled genomes (MAGs) were also included. Approximately 3,100 clone sequences (47 new) are present in AcetoBase clone dataset. In AcetoBase version 1, the taxonomy prediction for the clone sequences was done with SINTAX algorithm (Edgar, 2016). However, in version 2, the taxonomy for the complete clone dataset was predicted by acetotax program of AcetoScan pipeline (Singh et al., 2020) using AcetoBase reference protein (version 2) dataset, accessible at https://acetobase.molbio.slu.se/download/ref/1. Acetotax program is an unsupervised sequence annotation program which filters out non-FTHFS sequences, performs best open reading frame (ORF) analysis and annotate taxonomy to FTHFS sequences based on latest version of AcetoBase. The new taxonomic prediction of clone sequences (best ORF) indicated that most clones generated for FTHFS sequences belong to three major phyla *i*.*e*., Firmicutes (2355/3061 sequences, 77%), Proteobacteria and Spirochaetes. The taxonomic coverage of clone sequences present in AcetoBase is presented in **Fig. 2**. For the ease of visualization, sequences (2355) associated with phylum Firmicutes were not included in **Fig**. The taxonomy prediction for the clone sequences helped in correctly associating the clone sequences to a bacterial species. For instances among the sequences generated in a study by Parameswaran et al., 2010, none of the sequences submitted as uncultured *Alkaliphilus* sp. clone (10 clone sequences) belong to *Alkaliphilus*, in fact 46/47 clone sequences available from this study were more closely related to *Acetobacterium* spp. (>94% blastx similarity) (**Fig. 4**). These clone sequences with putative taxonomy and species level percentage identity are available in AcetoBase clone database with accession number CN_0000003015-CN_0000003061. Furthermore, the phylogenetic trees for the reference protein, nucleotide and clone datasets were also reconstructed according to the updated database content and information about phylogenetic tree construction is accompanied at the web-interface.

**Figure 1:**
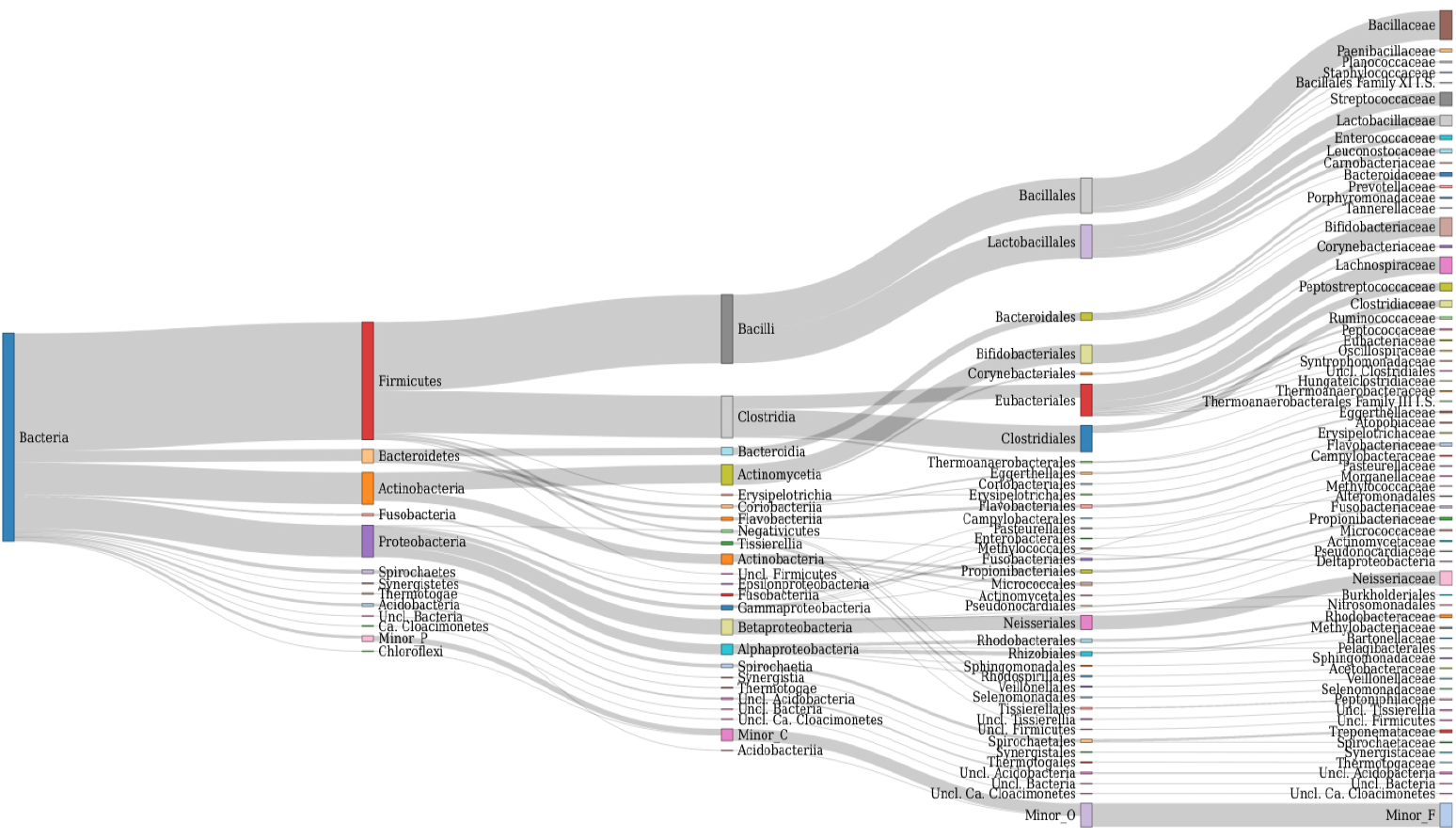
Taxonomic coverage of FTHFS sequences present in AcetoBase up to the family level. The candidates with less than 50 sequences are merged in minor taxa.

**Figure 2:**
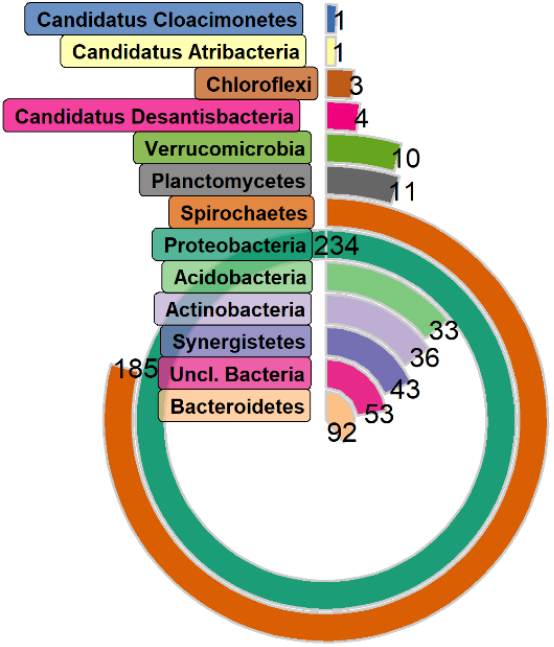
Taxonomic coverage of clone sequence dataset of AcetoBase at the phylum level except Firmicutes (2355). The number in the figure represents the sequence count belonging to respective phyla.

### RibocetoBase dataset

Although in AcetoBase version 1, FTHFS sequences were connected to 16S rRNA genes for the respective bacteria, these sequences were not available as a reference dataset for taxonomic annotations. In version 2, we have included RibocetoBase which can be readily downloaded and used for the taxonomic annotations of FTHFS containing community from 16S rRNA gene sequences. RibocetoBase is a 16S ribosomal RNA gene sequence dataset of the FTHFS harbouring bacterial community and it contains around 9,169 sequences. The details for the RibocetoBase and its development can be found elsewhere (Singh et al., 2021b). The taxonomic coverage of the five major (>100 sequences, 8,733/9,169) and 13 minor (10-100 sequences, 401/9,169) phyla are presented in **Fig. 3A** and **3B**, respectively. Around 20 Phyla, having less than 10 sequences (35/9,169), are ignored in the visualization. RibocetoBase taxonomic lineages are according to the genome taxonomy database (GTDB) (Chaumeil et al., 2019). RibocetoBase dataset can be accessed at https://acetobase.molbio.slu.se/download/ref/3.

**Figure 3:**
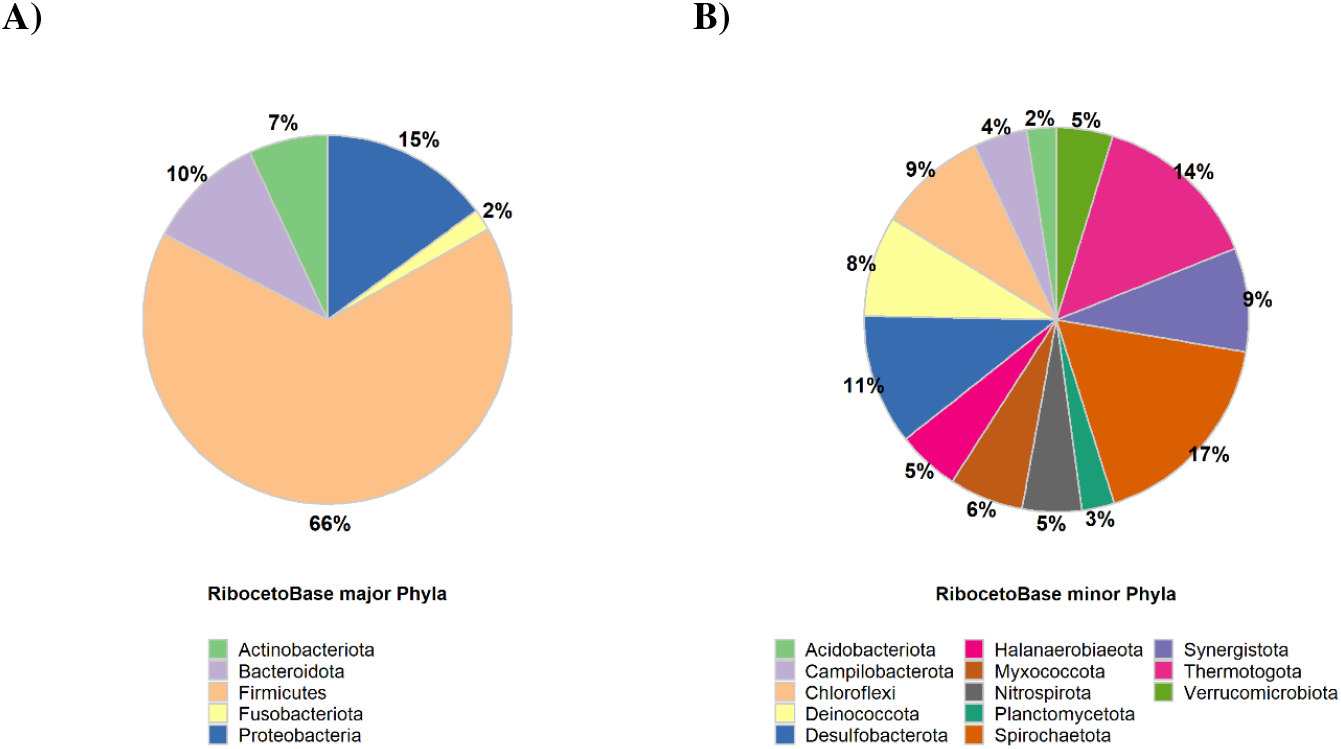
Taxonomic distribution of RibocetoBase dataset of FTHFS harbouring communities present in AcetoBase at the phylum level. **A)** represents the RibocetoBase major phyla having >100 sequences and **B)** represents RibocetoBase minor phyla with sequence count 10-100.

### Enhanced web-interface and functionalities

FTHFS has been used as a marker gene for acetogens and several acetogens have been isolated and characterized over the years. However, these was a lack of an elaborate collection of the original articles and micrographs in which these acetogens were isolated, characterised and described. In AcetoBase version 2, we have included a collection of micrographs of acetogens, and links has been provided to the original articles describing the isolation and description of the acetogen. In case the original article mentioning the isolation and characterization of an acetogen do not have a micrograph, images from other publicly available articles have been retrieved. A check mark represents the original article associated with respective acetogen. Additionally, we have also linked the acetogen to NCBI taxonomy database which would be helpful in 1) getting complete information of different strains of respective species 2) taxonomic names and synonyms and its state of validation 3) linking out to other NCBI databases *via* NCBI taxonomy database for extra information. AcetoBase information resource for acetogens is accessible at https://acetobase.molbio.slu.se/organism/acetogen.

As it is well known and mentioned earlier, FTHFS, although a marker for acetogens, is also present is some syntrophic acid degrading bacteria and sulphate reducing bacteria (Lovell, 1994; Singh et al., 2019, 2020). In several previous studies on FTHFS gene amplicon-based community profiling, for instance studies by Leaphart and Lovell, 2001; Leaphart et al., 2003; Lovell and Leaphart, 2005; McSweeney et al., 2009; Singh et al., 2021b, 2021a *etc*. syntrophic acid degrading bacteria and sulphate reducing bacteria have been reported together with acetogenic bacteria. These group of bacteria uses FTHFS gene, for instance, in biosynthesis of folate or short chain fatty-acid oxidation and is not involved in reductive acetogenesis like acetogens. Thus, considering the prospects and potential of FTHFS gene profiling in different environments, we have included information (similar to acetogen as described above) of FTHFS harbouring syntrophic acid degrading and sulphate reducing bacteria. Worth noting is that not all syntrophic acid degrading bacteria and sulphate reducing bacteria possess FTHFS gene in their genome and AcetoBase only includes the bacterial species which harbours FTHFS gene. We have also grouped these bacteria according to the functional categories as they have been described *viz*. syntrophic acetate oxidising bacteria (SAOB), syntrophic propionate oxidising bacteria (SPOB), syntrophic butyrate oxidising bacteria (SBOB), syntrophic fatty acids oxidising bacteria (SFOB), sulphate reducing bacteria (SRB) and syntrophic benzene oxidising bacteria (SBzOB). This systematic grouping and collection of syntrophic bacteria can be reached at https://acetobase.molbio.slu.se/organism/syntroph.

### Search functionality and results

The new version of AcetoBase now supports search with query from any taxonomy level (phylum-strain) and regular expressions (matching word patterns) in the search bar. The search results are rendered with the link out functionality to respective NCBI database for genome and protein identifiers. The complete lineage for the search results is displayed with link out to NCBI taxonomy database from the species name (**Fig. 4**). The strain level information is available for the bacterial species and MAGs wherever possible. With respect to the clone dataset, search now also supports query with taxa name, regular expressions and search with isolation source variables, for example, anaerobic digester, gut, roots, colon, microbial fuel cell *etc* (**Fig. 4**). The search results for the clone dataset now also accompanies the putative taxonomy predicted for the clone sequences, as mentioned earlier. The species level percentage identity is available with the lineage for each clone identifiers (**Fig. 4**).

**Figure 4:**
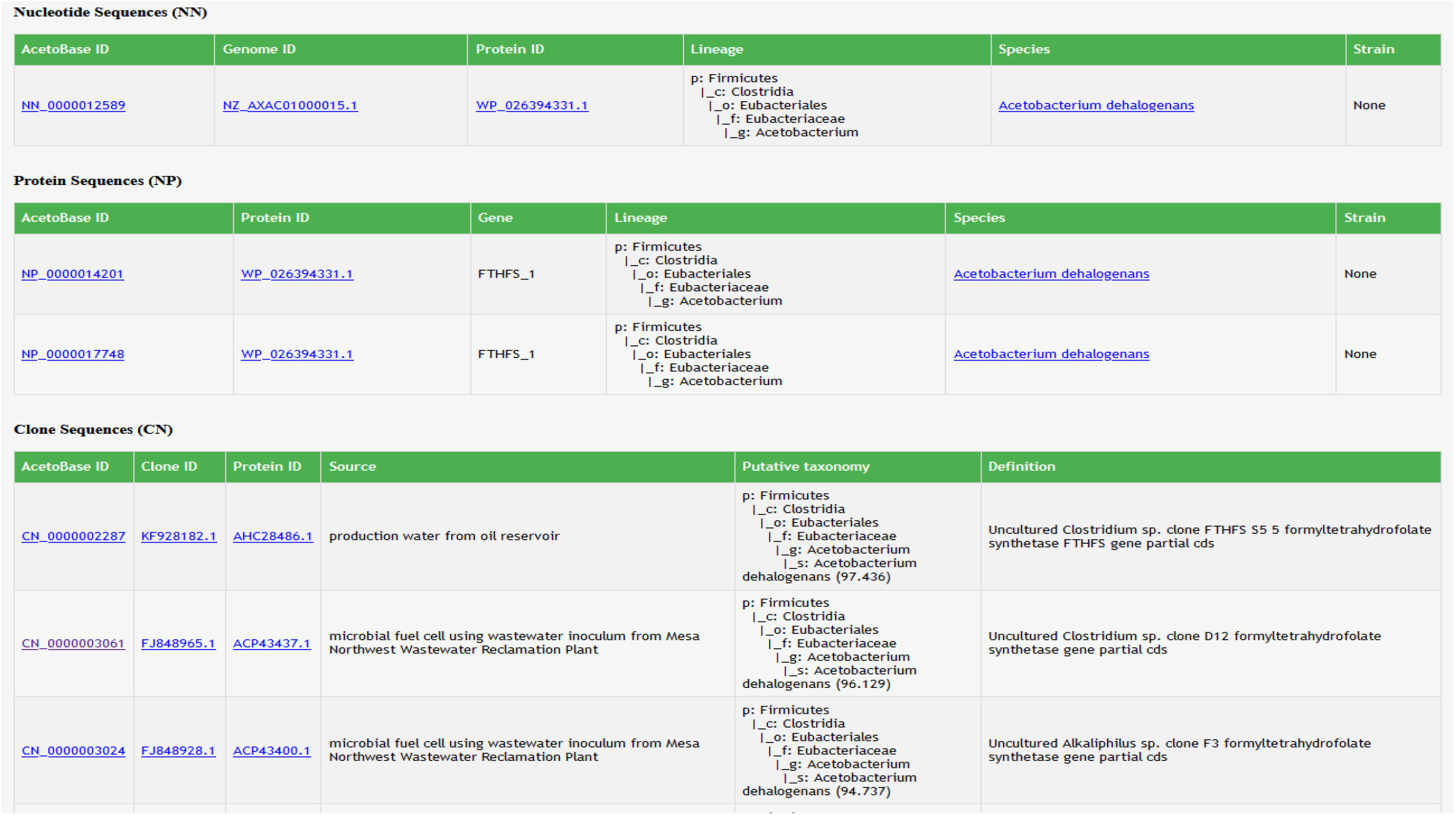
A screenshot of the AcetoBase version 2 search results showing taxonomic lineages of the query against the nucleotide and protein datasets. The search results for the clone dataset illustrates the isolation source of the clone, its description/definition and the putative taxonomy predicted with the acetotax program.

### REANALYSIS OF FTHFS HIGH-THROUGHPUT SEQUENCING DATA

AcetoBase (version 1) was published in 2019 and has been used for the high throughput sequencing data-analysis in our previous studies (Singh et al., 2020, 2021b, 2021a). Thus, to evaluate the impact of the AcetoBase update on the taxonomic annotation of the FTHFS OTUs generated by AcetoScan pipeline, we reanalysed the data from respective studies using AcetoBase version 2. The methodology of the reanalysis is described in the following.

### Methodology

The reanalysis were done for the data associated with the articles Singh et al., 2020 (dataset 1), Singh et al., 2021b (dataset 2) and Singh et al., 2021a (dataset 3), respectively. The reanalysis of these datasets (separately) was done with AcetoScan pipeline using the parameters described in the respective papers and AcetoBase protein dataset (version 2) as a reference database. In the present study the results from reanalysis of the respective dataset were compared with results from the previous analyses based on the differences and variations in the taxonomic annotation. Thus, for the ease of understanding in the following text, the expression “previous study/analysis” should be considered as the results published for respective dataset/paper and word “reanalysis” should be considered as the reanalysis of respective dataset in present study. The specific detail of the reanalysis is described, and major community insights were discussed in the following:

### Reanalysis of dataset 1

In the present paper for the reanalysis of dataset 1, we have in addition to the previous study also included additional new set of FTHFS sequence data for reactor GR1, which was not presented in study by (Singh et al., 2020) (**Supp. Fig. 1A-F**). The analysis in the original article was performed separately for the forward and reverse reads. The reanalysis in present article for both reactor GR1 and GR2 was conducted by merging the forward and reverse reads of each respective samples. The analysis was performed with 100% clustering threshold and using parameters -m 300 -n 120 -q 20 -c 5 -t 1.0. The previous study described 1,171 and 1,241 OTUs obtained at 100% clustering from forward and reverse reads in reactor GR2, respectively. The reanalysis of dataset 1 and new data from the reactor GR1 resulted in 1,024 OTUs at 100% clustering threshold while using merged data from forward and reverse reads. The results from reanalysis indicated towards higher percentage of annotated sequences as compared to the previous analysis and with no unclassified operational taxonomic units (OTUs) at genus level (**Supp. Fig 1E**). The most distinctive change was observed for the classification of the OTUs from genus *Flavonifractor* to unclassified Lachnospiraceae. Genera related to OTUs which were not classified in present analysis but were classified in earlier analysis were *Butyrivibrio, Caldisalinibacter, Eggerthella, Hungateiclostridium, Lagierella, Maledivibacter* and *Phocea*. Some taxa were classified in the present analysis, which were not classified in previous study, *e*.*g*., *Peptoniphilus, Ruminiclostridium, Sedimentibacter, Thermoanaerobacter* and *Urinicoccus*. The genus *Senegalimassilia* was only observed in the new additional data from reactor GR1 which was not used in original research where only reactor GR2 was used (**Supp. Fig 1E**).

### Reanalysis of dataset 2

The analysis of dataset 2 was done for forward reads as described in the original article with the analysis parameters -m 300 -n 150 -q 21 -c 10 -r 1, -e 1e-30 -t 1.0. The number of OTUs (387) generated in the present study was similar to the number of OTUs generated in previous study (391). The reanalysis results (**Supp. Fig 2A-F**) from dataset 2 at phylum level showed significant reduction in the annotation of the OTUs which were classified earlier as Actinobacteria and in contrast to previous analysis no annotations were obtained for Spirochaetes at relative abundance (RA) greater than 1% (**Supp. Fig 2A**). Compared to the previous study, the fraction of OTUs classified as Firmicutes increased. At the genus level, the genera which were not classified in the reanalysis were *Caloranaerobacter, Marvinbryantia, Methylocystis, Oscillibacter, Phocea, Tissierella, Treponema, Varibaculum* and unannotated (NA) OTUs (**Supp. Fig 2E**). Clear and significant differences in the taxonomic annotations were noted for the OTUs previously annotated as *Marvinbryantia* and *Varibaculum*, which were in the new analyses annotated as unclassified Lachnospiraceae and *Urinicoccus*. The total number of genera (RA >1%) annotated in reanalysis were 14 as compared to 21 in previous study. Following the changes at genus level, the most significant chance in the reanalysis was for the species belonging to phylum Cloacimonetes (**Supp. Fig 2F**). In current study, only one species was classified Candidatus Cloacimonetes bacterium (MAG ID: AS05jafATM_99) originated from anaerobic digesters (Campanaro et al., 2020) as compared to previously annotated two species Cloacimonetes bacterium HGW-Cloacimonetes-1 and Cloacimonetes bacterium HGW-Cloacimonetes-2, which were originated from a metagenomics project analysing subsurface sediments (Hernsdorf et al., 2017).

### Reanalysis of dataset 3

Dataset 3 were generated in a study by Singh et al., 2021a, in which a high-throughput microbiological surveillance of 6 different biogas plants/11 biogas reactors (1 plug flow and 5 parallel CSTR) were performed. The parameters used for the reanalysis were the same as used in previous study (-m 300 -n 150 -q 20 -c 5 -e 1e-30 -t 1.0) which resulted in 1909 OTUs, compared to 1901 OTUs in the previously analyses (**Supp. Fig. 3A-3F**). At the phylum level, the new analysis illustrated a similar community diversity in all reactors as previously presented, except for the reactor C6. The reanalysis results showed increase in the percentage of OTUs annotated as Firmicutes with corresponding decrease in OTUs for phylum Actinobacteria. Additionally, the OTUs previously belonging to Synergistetes were not observed in the reanalysis expect for one sample (**Supp. Fig 3A**). Due to the higher number of genera in both previous and present study, presentation and comparison of each genus and species is not included here was and the readers are advised to refer to **Supp. Fig 3E** and **3F** for comparison. However, the overview of annotation results indicates that annotation accuracy is significantly improved with lesser number of resulting taxa and with a higher percentage identity to the reference database as compared to previous study.

## Discussion

The present study reanalysed the FTHFS sequence data generated in previous studies using the updated versions of AcetoBase and compared the differences between taxonomic annotation. A careful interpretation of the reanalysis results from the present study, its comparison to previous respective studies (Singh et al., 2020, 2021b, 2021a) supports and approves the improvements in taxonomic annotations resulting from the updates in AcetoBase. Increase in the taxonomic diversity in reference database provides opportunity for the taxonomic annotations of OTUs with higher accuracy (greater percentage identity and lower e-value) in the best hit strategy used by AcetoScan. However, since FTHFS is not a taxonomic marker and due to technical limitations in FTHFS amplicons sequencing, the community diversity evaluated with FTHFS gene can have similarities but also differences if compared to 16S rRNA gene and whole genome metagenome-based community profiles. Still, a comparative analysis of FTHFS and 16S rRNA gene amplicons by Singh et al., 2021b demonstrated that the microbial community profiled with FTHFS showed high similarity to the structure and dynamics as revealed by community profile using 16S rRNA gene. The recent changes in the taxonomic lineages of bacteria have also contributed to the differences in the present and previous studies used in this paper.

## SCOPE OF ACETOBASE

Acetogenic bacterial community are metabolically remarkably diverse can are found in hugely varied environments like anaerobic digesters, animal gut and rumen, human gut, marine sediments, forest soil, peatlands, permafrost, electrochemical fuel cell *etc*. (Breznak and Kane, 1990; Kusel and Drake, 1994; Parameswaran et al., 2010; Zhang et al., 2011; Drake et al., 2013; Coolen and Orsi, 2015; Müller et al., 2016; Sagheddu et al., 2017; Martinez et al., 2019; Singh et al., 2021a, 2021b). In these environments, they are not only involved in the carbon cycling via acetate but also extensively involved in the metabolism of ethanol, methanol, formate, butyrate, lactate, vanillate, chlorinated compounds, cellulose, mono-tetra saccharides, methylated amines *etc*. (Wolin and Miller, 1994; Das and Ljungdahl, 2003; Lovell and Leaphart, 2005; Rey et al., 2010; Hügler and Sievert, 2011; Wen et al., 2015; Lechtenfeld et al., 2018; Yang, 2018; Picking et al., 2019; Kountz et al., 2020).

AcetoBase was initially developed and used for the acetogenic community analysis in anaerobic digestor/biogas environment. However, the taxonomic diversity and sequence database contains huge magnitude of bacterial species which are also present in different environments. As discussed earlier, FTHFS primers can target syntrophic acid oxidising, sulphate-reducing bacterial groups *etc*. but also different fermentative FTHFS harbouring bacteria. These fermentative bacteria plays important roles in various anaerobic environments and has in addition huge physiological importance for humans and animals *viz. Bacteroidetes, Bifidobacterium, Clostridium, Faecalibacterium, Lactobacillus etc*. (*e*.*g*. Collado et al., 2013; Panasevich et al., 2015; Induri et al., 2022). AcetoBase thus has immense potential to be a representative database for the FTHFS based microbial ecological analysis in varied environments. For instance, human gut harbours ample amount of acetogenic bacterial communities which are significantly involved in gut physiology even in the presence of methanogens and sulphate reducing bacteria (Doré et al., 1995; Rey et al., 2010). Further, members of acetogenic or FTHFS-possessing communities *viz. Clostridium, Blautia, Eubacteria, Eggerthella, Prevotella, Ruminococcus*, family Lachnospiraceae *etc*. are substantially involved in, for instance, dysbiosis of autoimmune disorders, multiple sclerosis, rheumatoid arthritis, systemic lupus erythematosus, cirrhosis, gastrointestinal disorder, Parkinson’s disease *etc*. (Floch, 2017; Punzalan and Qamar, 2017; Raghavendra and Pullaiah, 2018; de Oliveira, 2019; Keshavarzian et al., 2020). There has been only a few studies which studied the FTHFS harbouring communities in human gut (*e*.*g*. Ohashi et al., 2007, 2009), yet any large scale longitudinal studies on this subject is completely lacking.

## CONCLUSIONS

In this study, we present the updates made in AcetoBase in database content, functionality and user-interface and reanalysed the FTHFS sequence data published in our earlier studies. The update in the database content included addition of sequences from new bacterial species and MAGs. This increase in the taxonomic diversity was aimed to assist the FTHFS community profiling and taxonomic annotation with higher accuracy. The enhancements in the web-interface especially for the acetogens and syntrophic acid oxidising and sulphate reducing bacteria could serve as a knowledge bank for organisms which possess FTHFS gene. Further, the improved search functionality added will be helpful in the ease of query the database and knowing the taxonomic lineage of the respective bacteria. The putative taxonomy for the clone sequences is of significant help in determining the correct taxonomy (as described above), which can be different from the taxonomic associations of published clone sequences or for future FTHFS clone library-based studies.

The reanalysis of the FTHFS sequence data from our previous studies demonstrated and supported the usefulness of database update in improving the taxonomic annotations resulting from AcetoScan analysis. The variations in the taxonomic annotations using AcetoBase version 2 as compared to version 1 did not change the overall dynamics or interpretations of the community profiles. Majority of the changes in taxonomic annotations were observed at lower taxonomic levels and among members of same order or family. Addition of new sequences from MAGs helped in better identification of FTHFS sequences. As amplicon sequence analysis is dependent on reference databases, continuous updates in databases are necessary and further addition of new sequences in future may still help in improving the taxonomic annotation. As AcetoBase contains sequences from acetogens and syntrophic organisms as mentioned above, detection of this acetogenic and syntrophic composite employing FTHFS amplicon sequencing could open new avenues in ecology and functional studies of microbial interactions in different environments especially anaerobic bioprocesses and gut of animal and humans. The ecological studies and understanding regarding syntrophic microbial interactions are scarce and longitudinal FTHFS profiling and faster analyses of community dynamics could be an outstanding tool in getting insights in the unknown syntrophic microcosm.

## DATA AVAILABILITY STATEMENT

The FTHFS raw sequence data from the reactor GR1 analysed in this study and associated to previous study (Singh et al., 2020) have been submitted to NCBI SRA (study: SRP336508) with BioProject accession number PRJNA761914. The OTU sequences generated by the reanalysis of dataset 2 and 3 have been submitted to AcetoBase with accession number UN_0000023501**-**UN_0000023887 and UN_0000023888-UN_0000025796, respectively.

## AUTHOR CONTRIBUTIONS

**Abhijeet Singh:** Conceptualization, Data Curation, Methodology, Software, Visualization, Writing - Original Draft. **Anna Schnurer:** Conceptualization, Funding acquisition, Resources, Writing - Review & Editing.

## ACKNOWLEDGEMENT

We would like to thank Hans-Henrik Fuxelius for his support and help in the original code modification and developing the version 2 of AcetoBase.

## SUPPLEMENTARY MATERIAL

Supplementary data 1, 2 and 3

## Supplementary Figures: 1A-F

### Description

The supplementary figures 1A-F (phylum-species level) are generated in present study by the reanalysis of the FTHFS amplicon sequence data (reactor GR2) from the study by Singh et al., 2020 and additional data mentioned in the present study (reactor GR1).

---

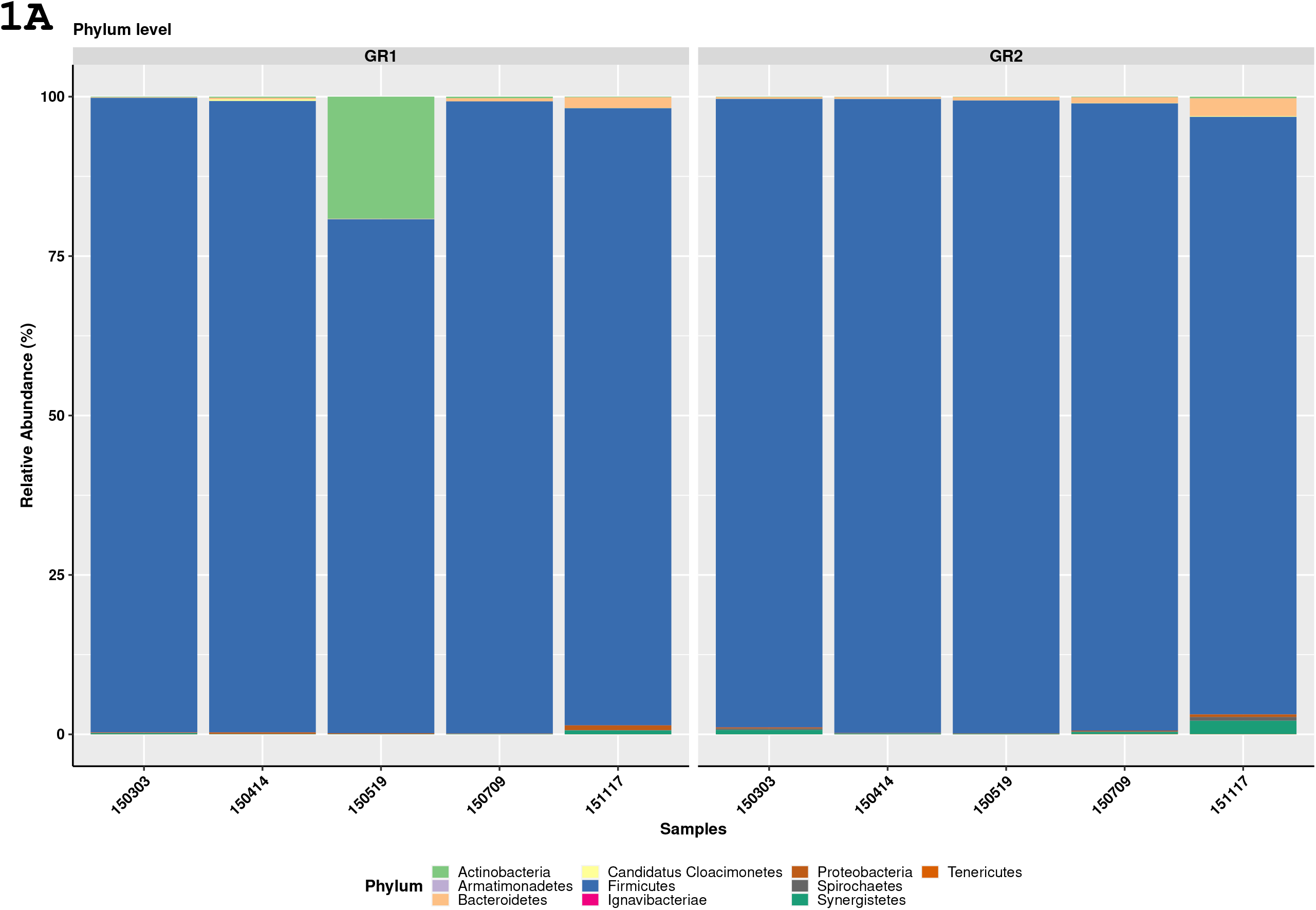
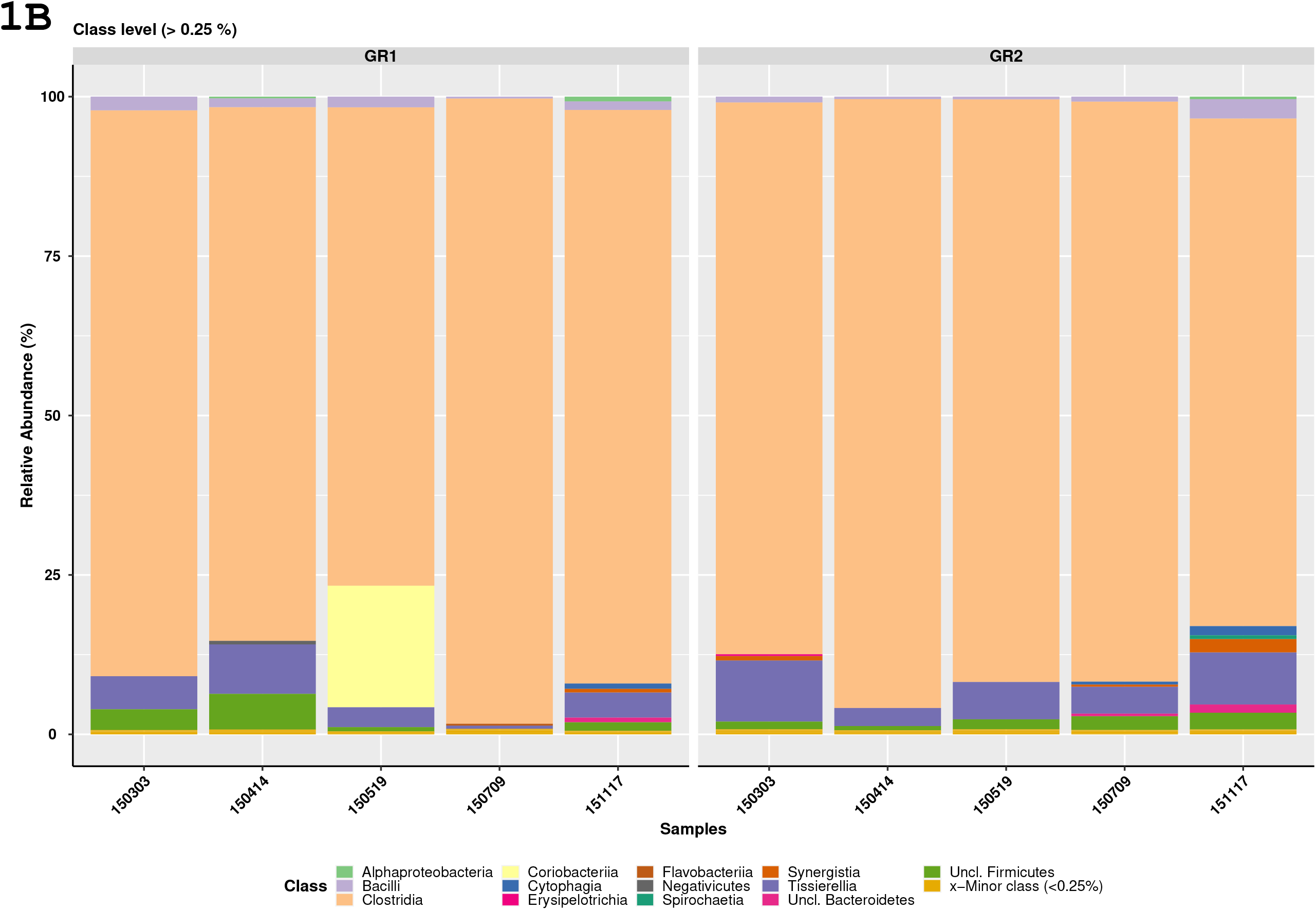
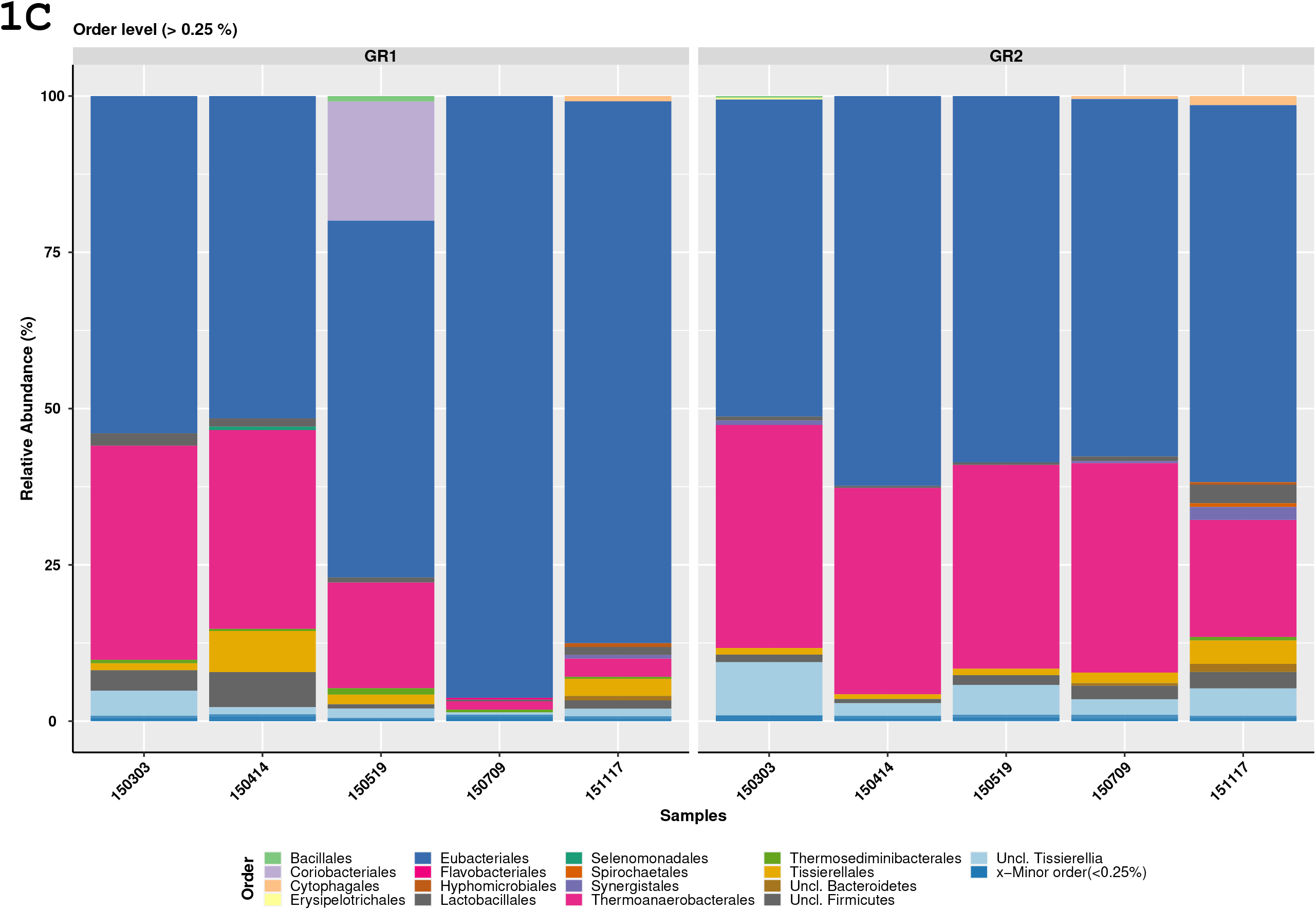
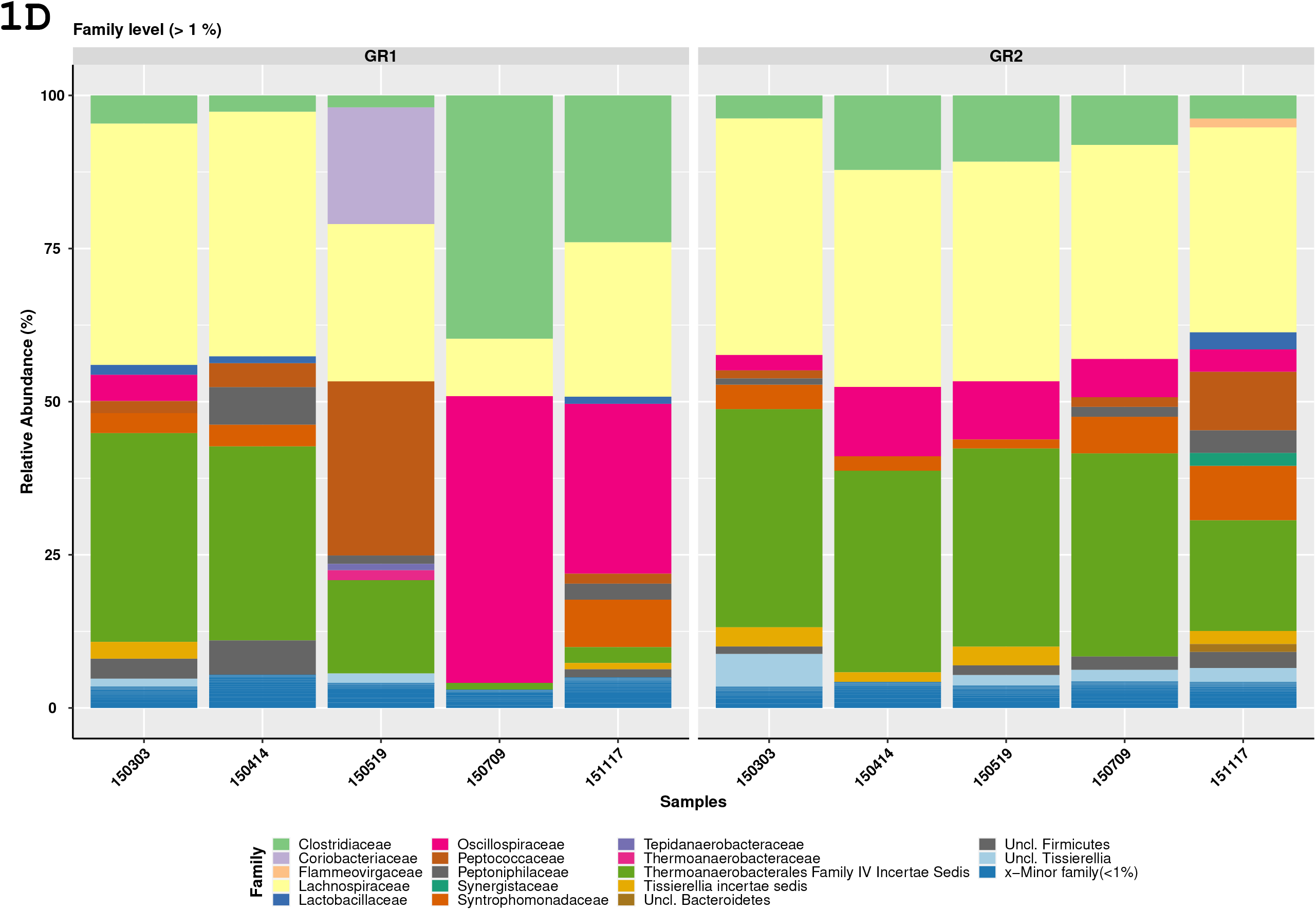
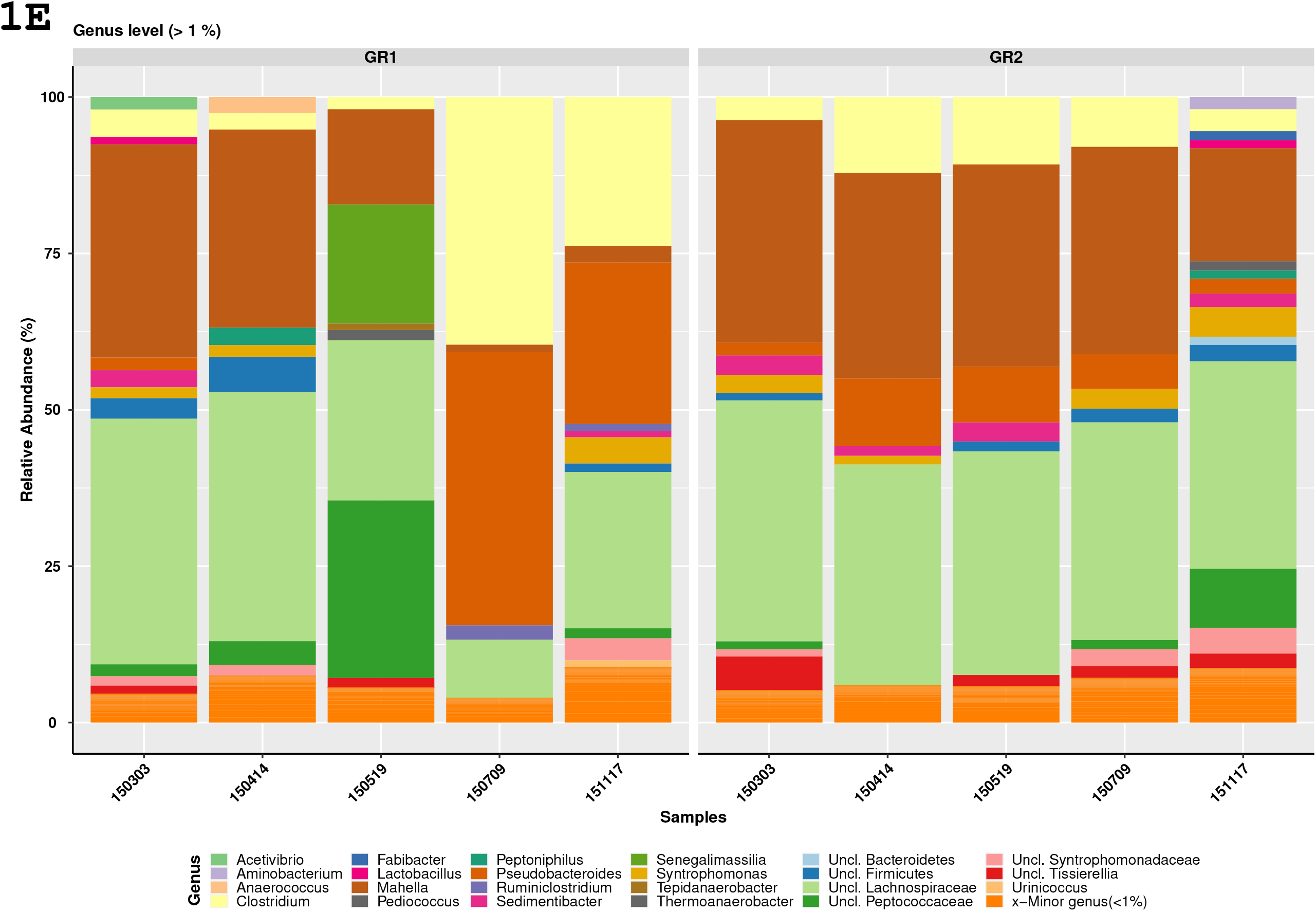
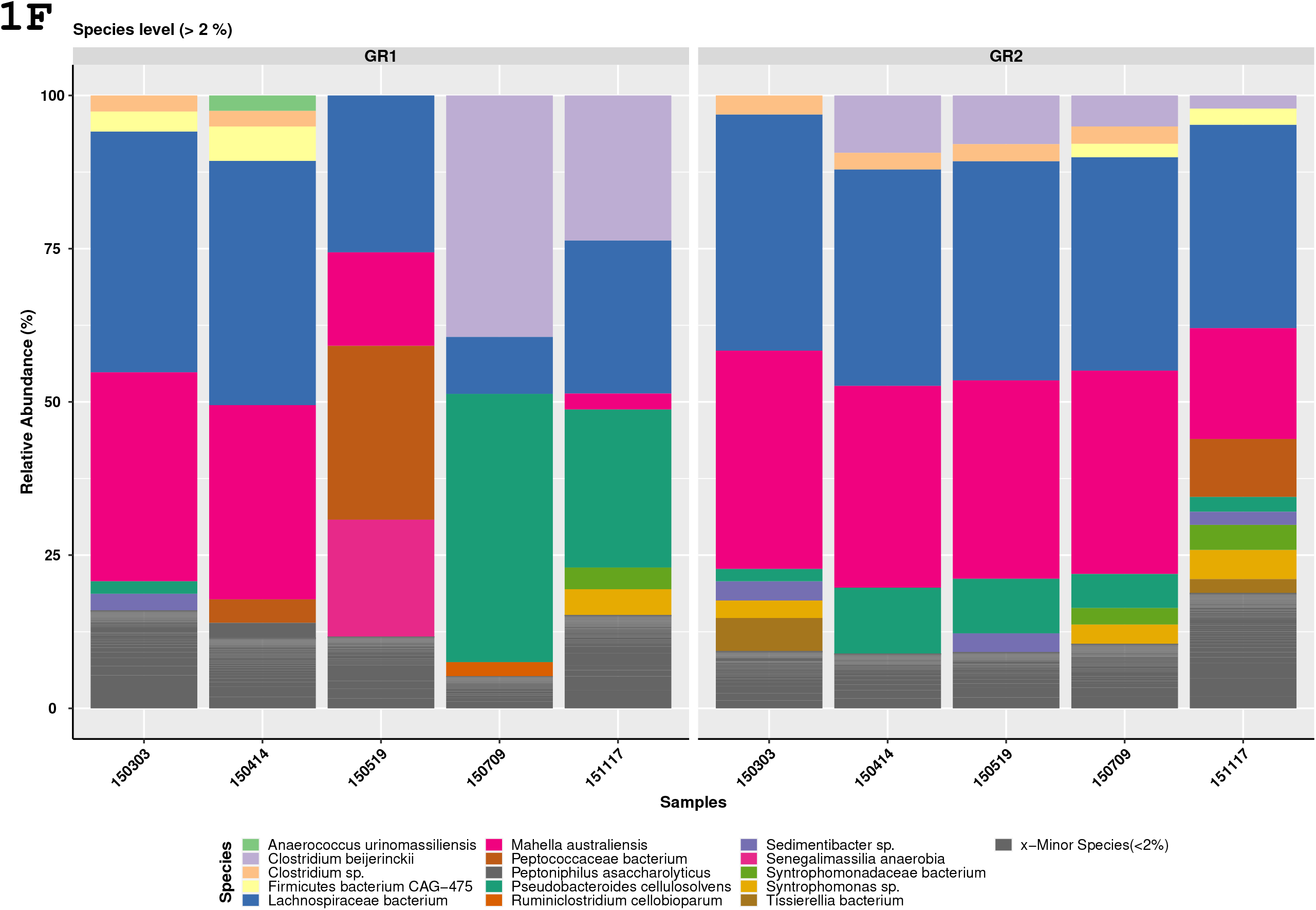

## Supplementary Figures: 2A-F

### Description

The supplementary figures 2A-F (phylum-species level) are generated in present study by the reanalysis of the FTHFS amplicon sequence data from the study by Singh et al., 2021.

---

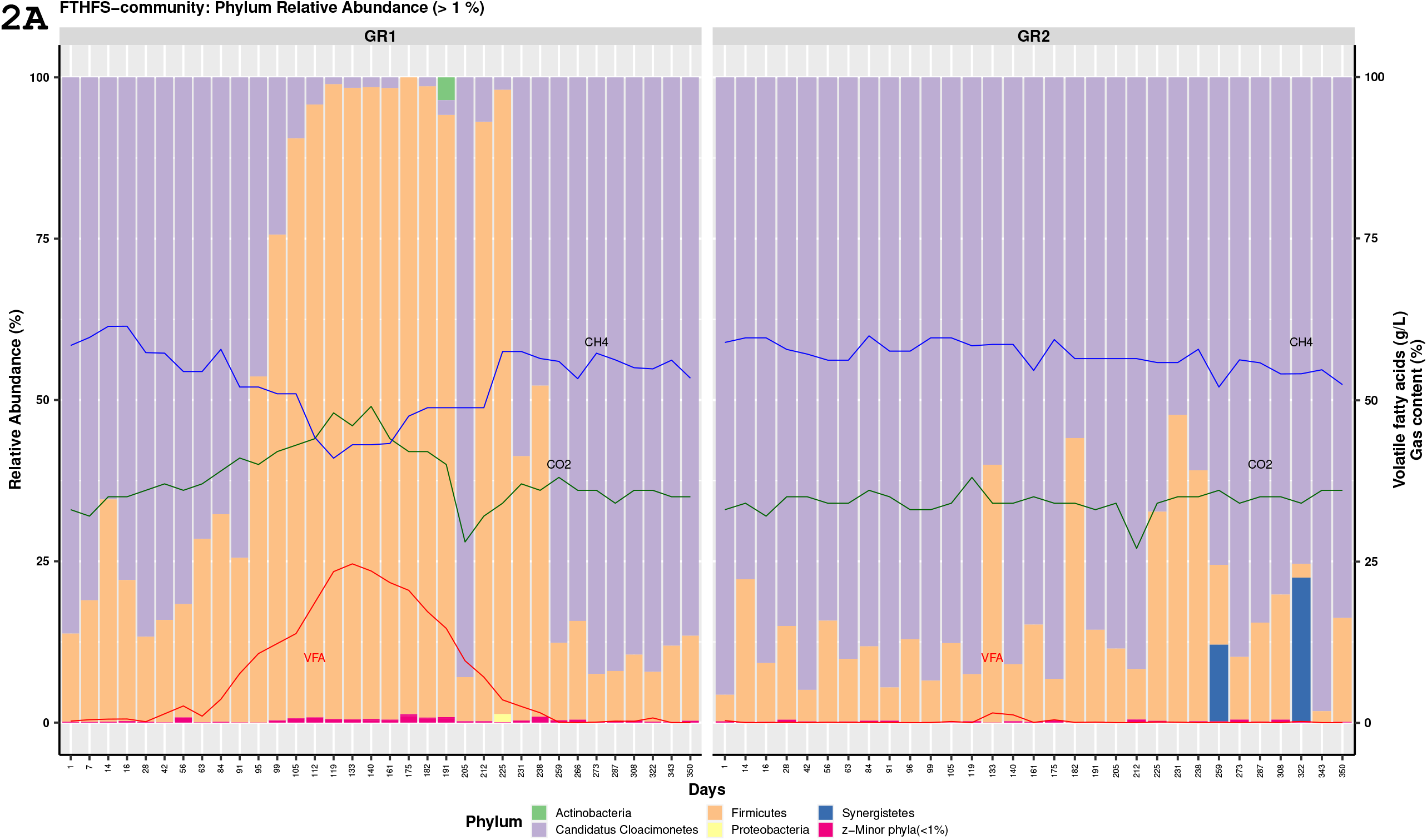
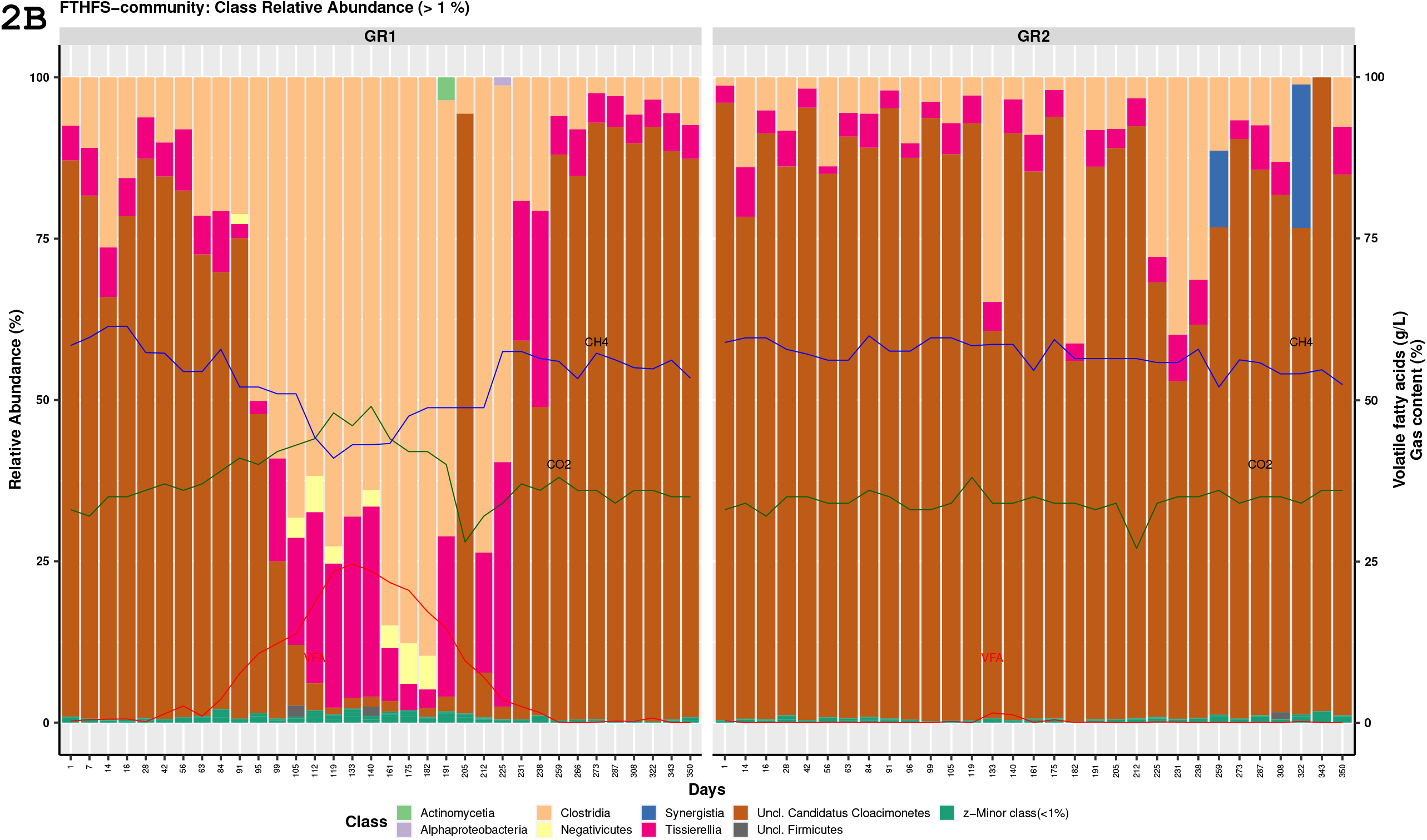
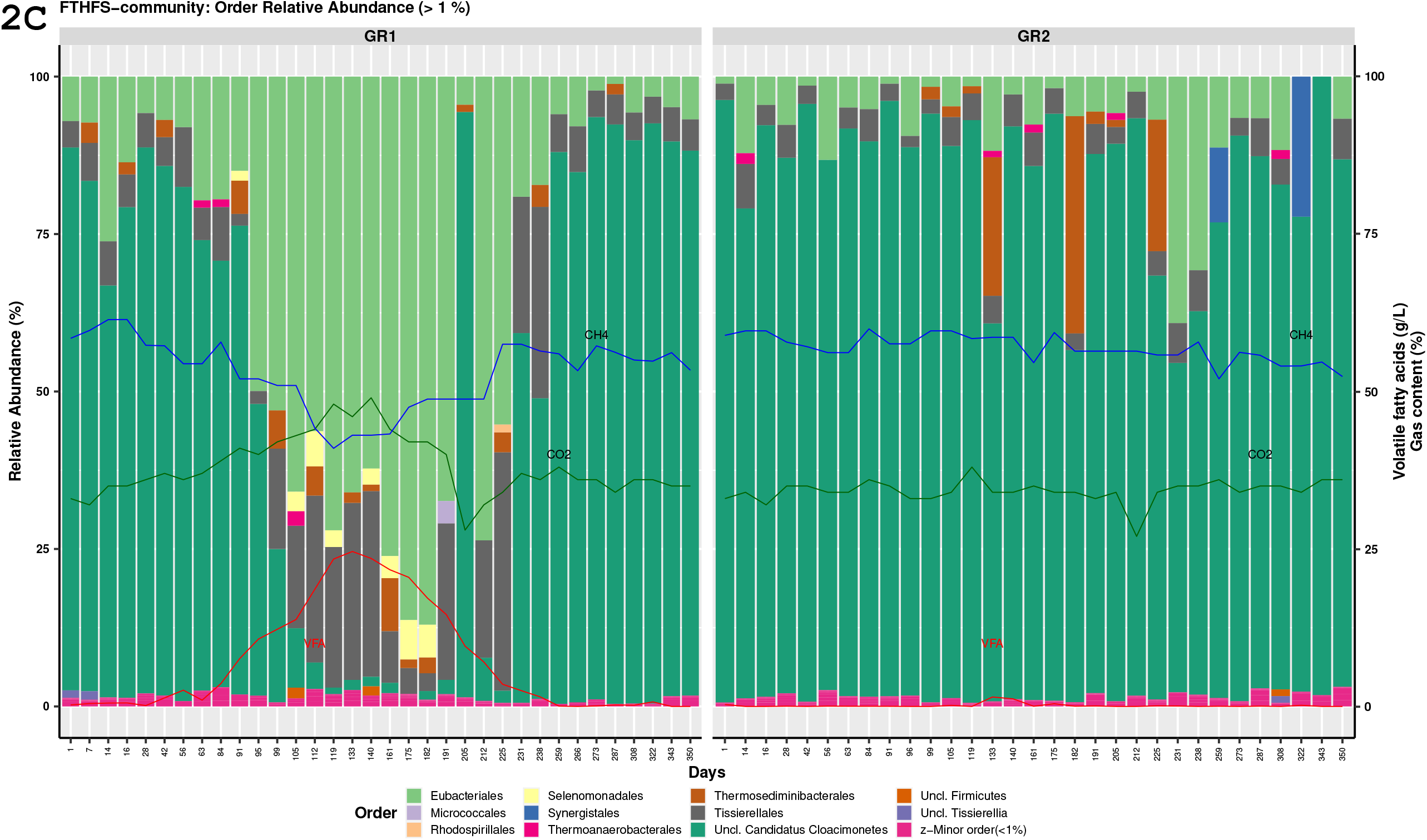
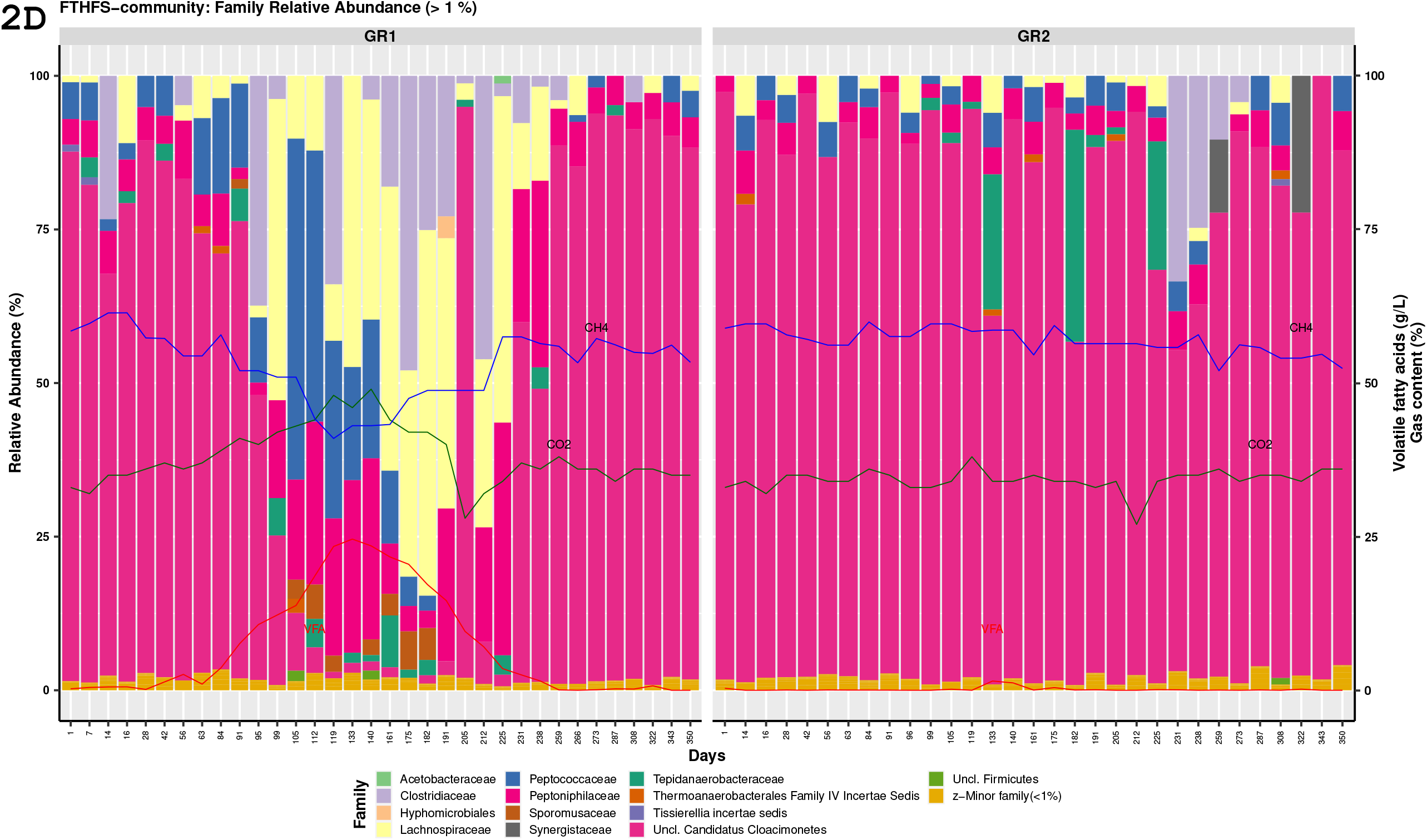
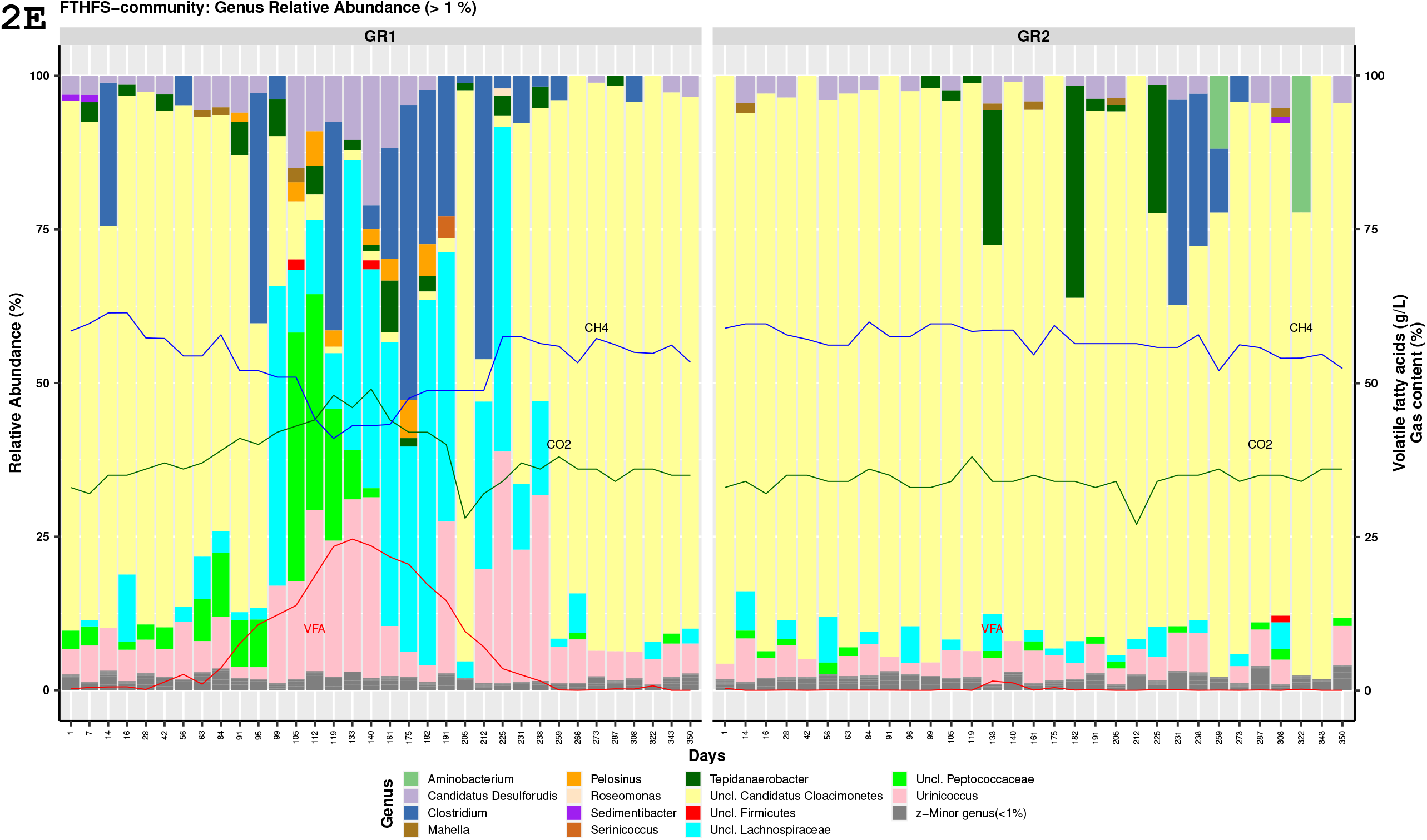
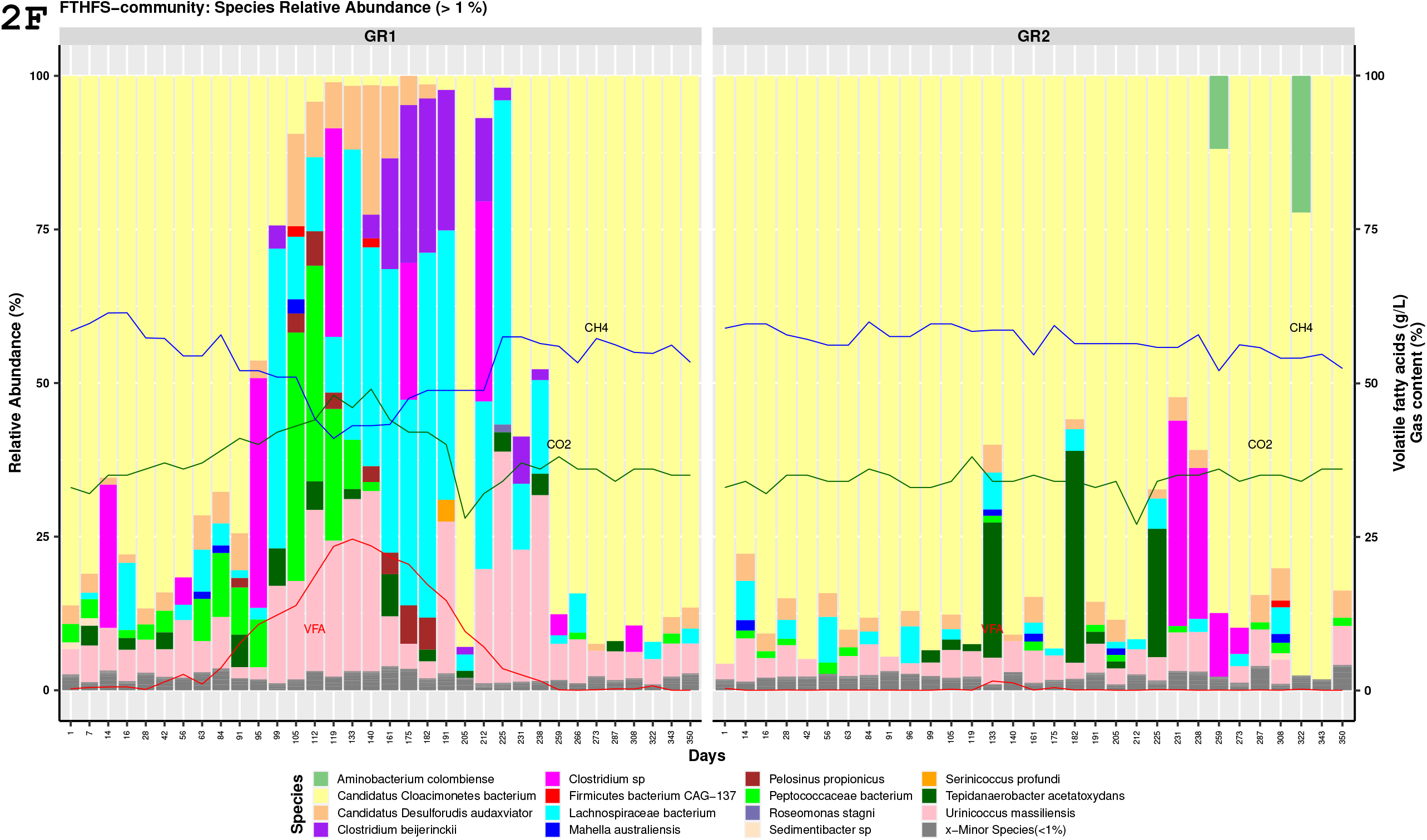

## Supplementary Figures: 3A-F

### Description

The supplementary figures 3A-F (phylum-species level) are generated in present study by the reanalysis of the FTHFS amplicon sequence data from the study by (Singh et al., 2021).

---

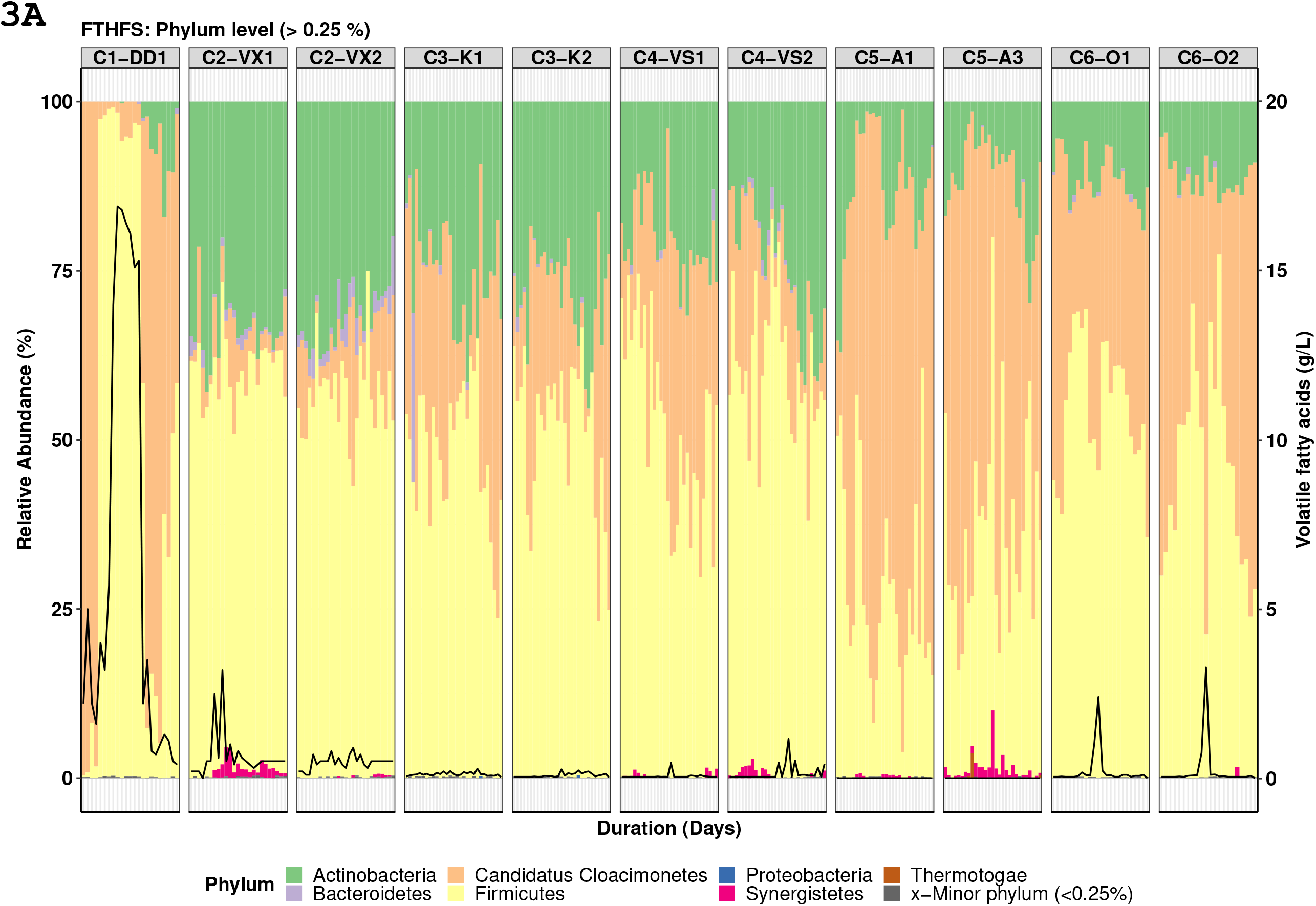
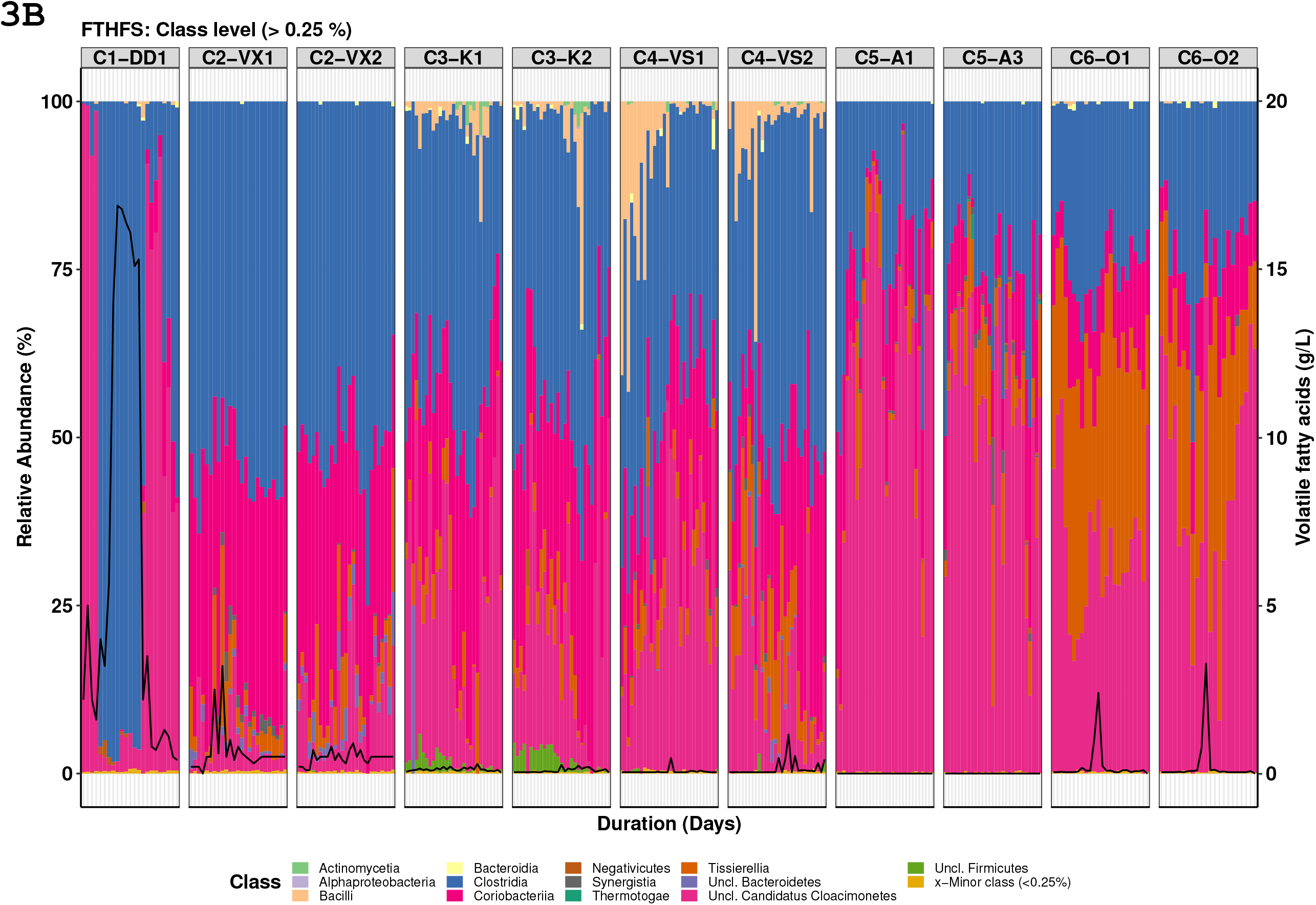
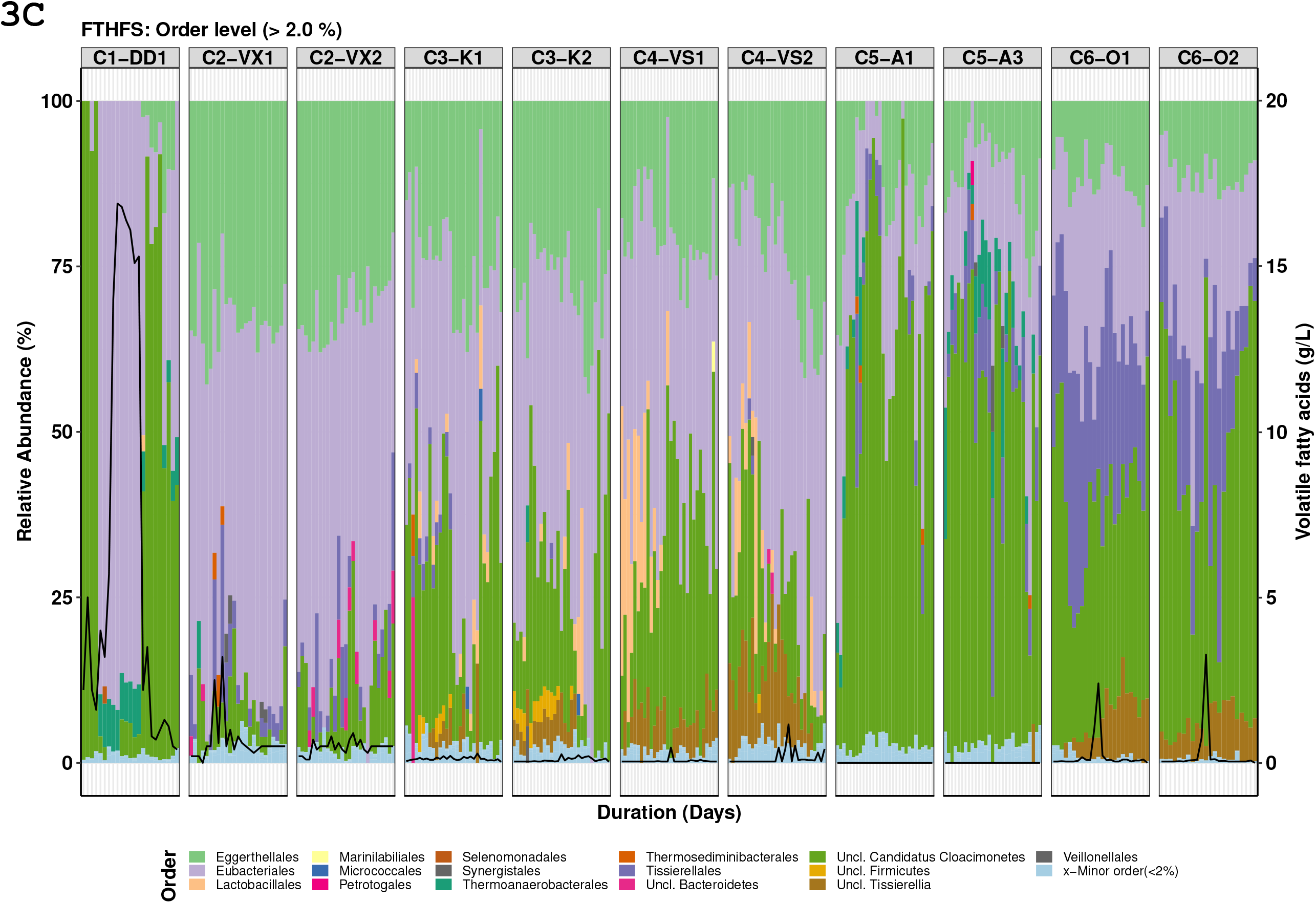
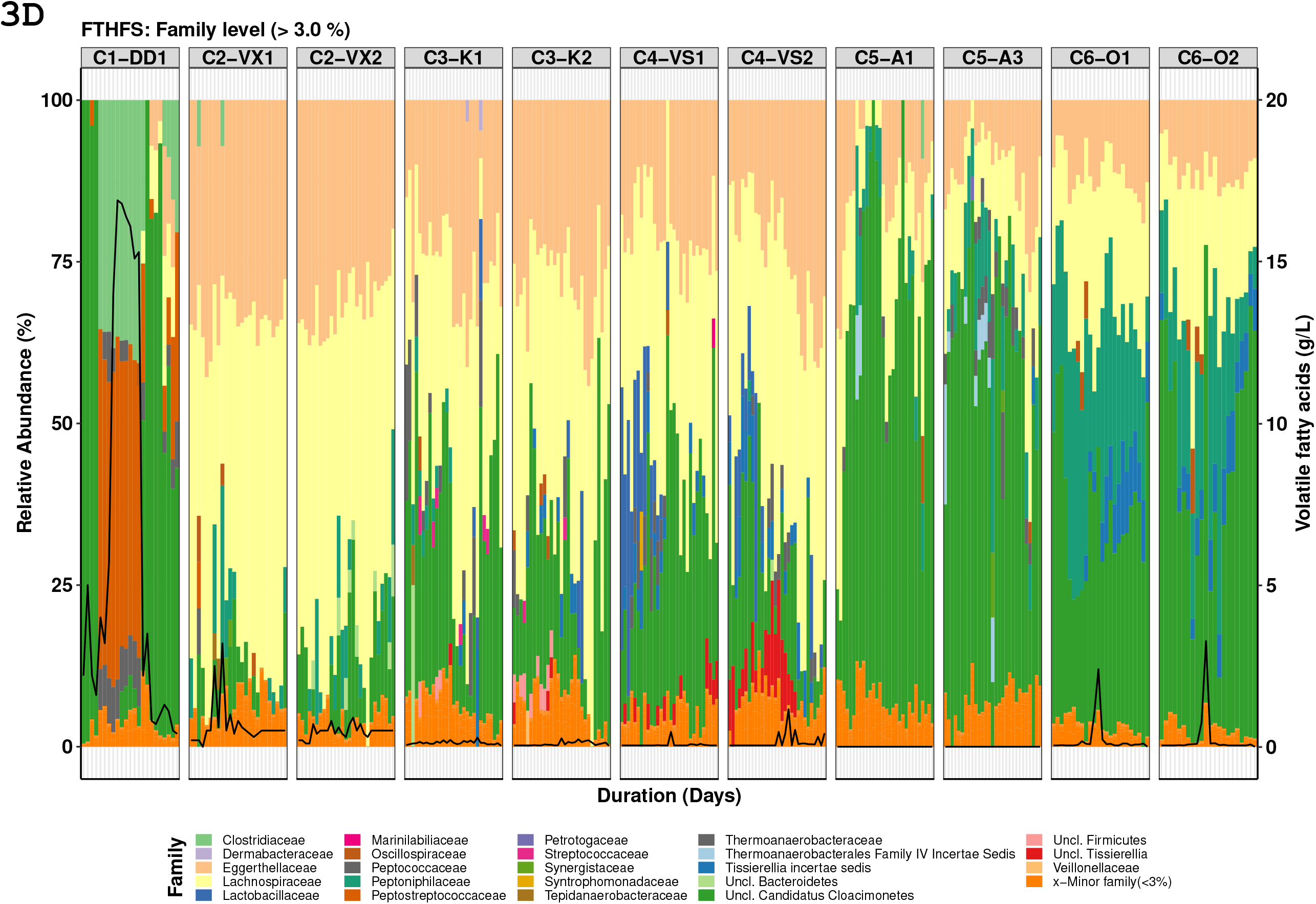
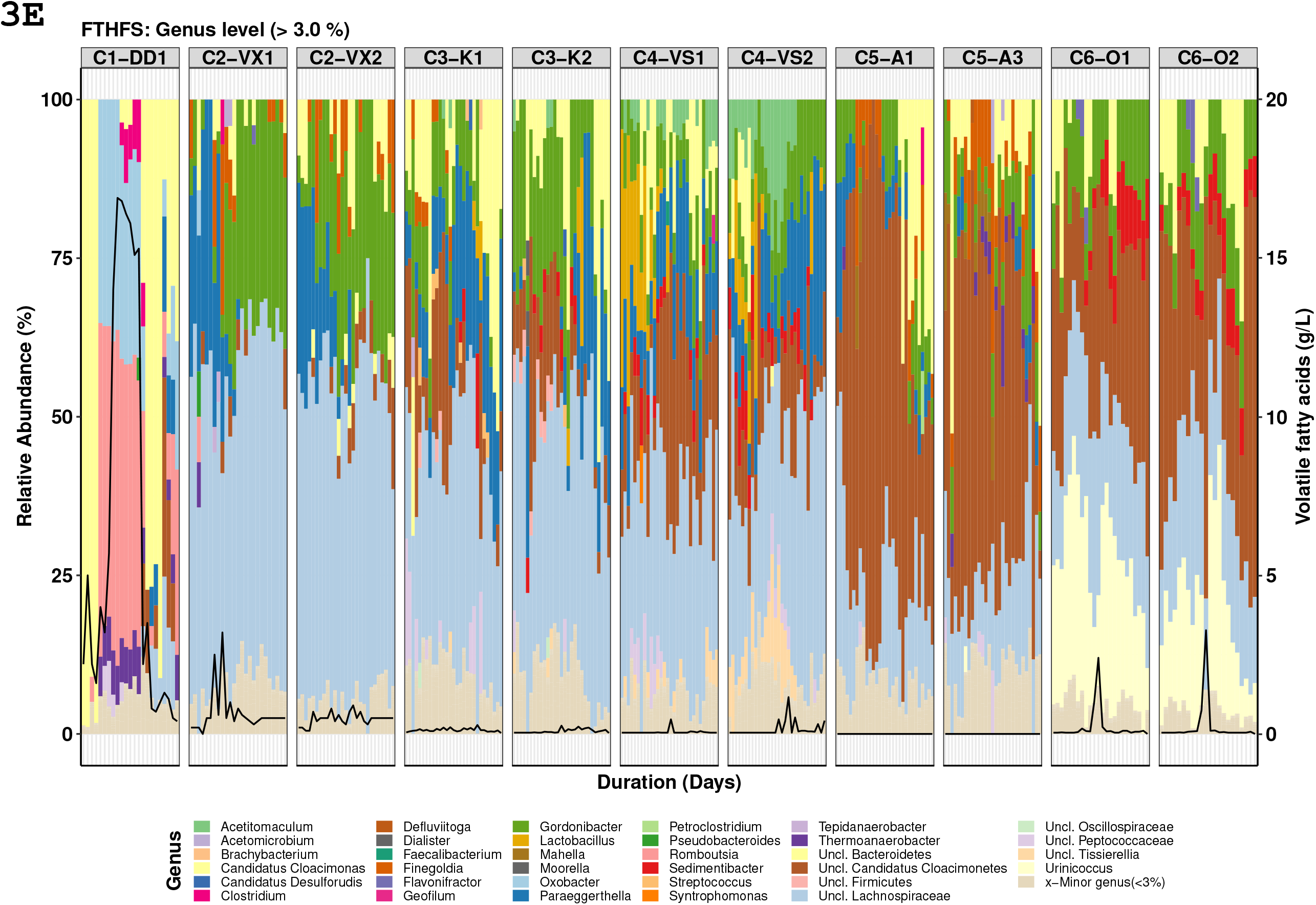
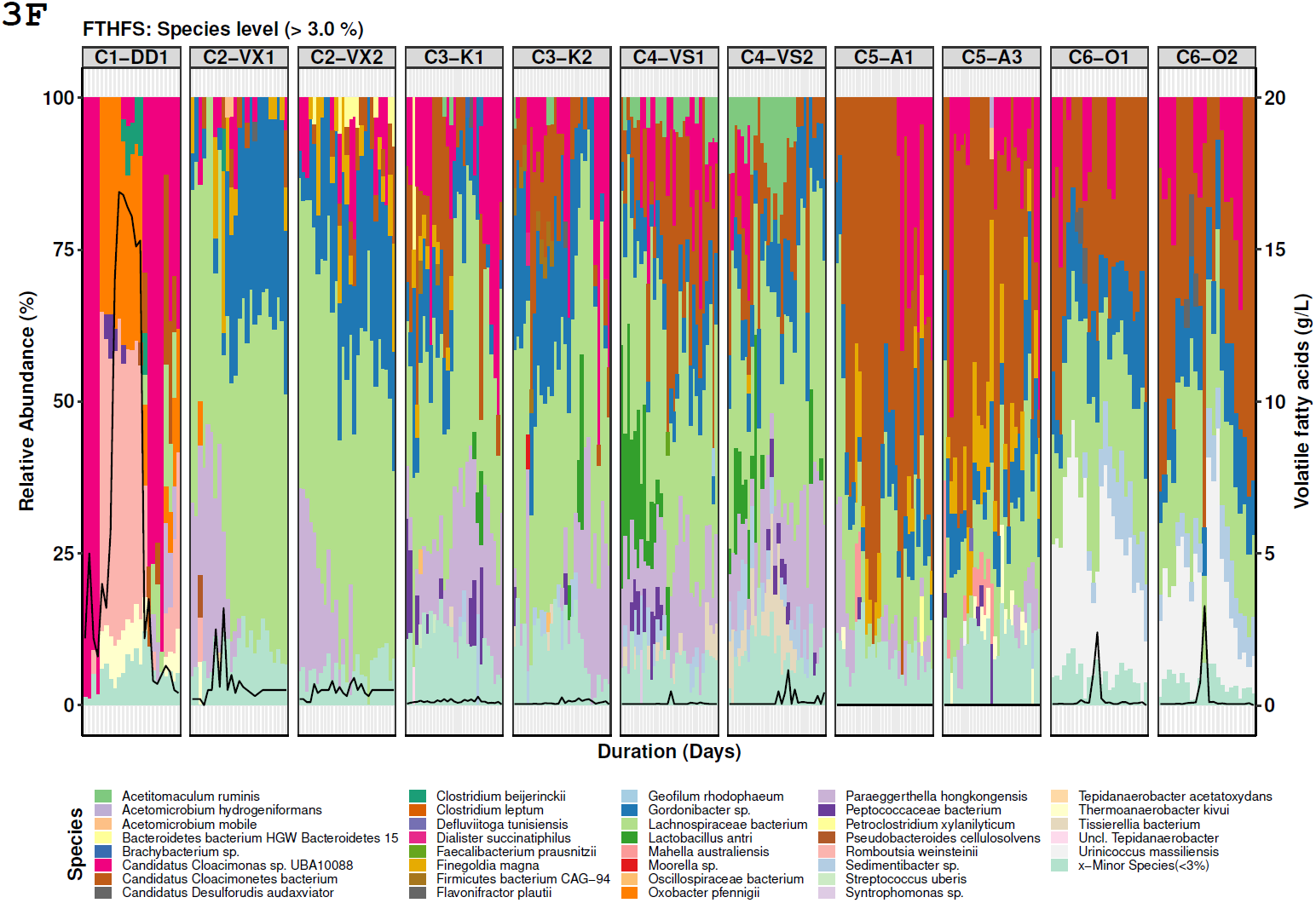

